# Data-driven approaches for Tau-PET imaging biomarkers in Alzheimer’s disease

**DOI:** 10.1101/244574

**Authors:** Jacob W. Vogel, Niklas Mattsson, Yasser Iturria-Medina, T. Olof Strandberg, Michael Schöll, Christian Dansereau, Sylvia Villeneuve, Wiesje M. van der Flier, Philip Scheltens, Pierre Bellec, Alan C. Evans, Oskar Hansson, Rik Ossenkoppele, the Alzheimer’s Disease Neuroimaging Initiative & the Swedish BioFINDER study

**Affiliations:** Montreal Neurological Institute, McGill University, Montreal, QC, Canada; Alzheimer Center and Department of Neurology, VU University medical center, Amsterdam Neuroscience, Amsterdam, Netherlands; Clinical Memory Research Unit, Lund University, Lund, Sweden; Memory Clinic, Skåne University Hospital, Lund, Sweden; Department of Neurology, Skåne University Hospital, Lund, Sweden; Wallenberg Centre for Molecular and Translational Medicine, University of Gothenburg, Gothenburg, Sweden; Department of Computer Science and Operations research, Université de Montréal, Montreal, QC, Canada; Centre de Recherche de 1’Institut Universitaire de Gériatrie de Montréal, University of Montreal, Montreal, QC, Canada; Department of Psychiatry, McGill University, Montreal, QC, Canada; Department of Epidemiology and Biostatistics, VU University medical center, Amsterdam, Netherlands

## Abstract

Previous positron emission tomography (PET) studies have quantified filamentous tau pathology using regions-of-interest (ROIs) based on observations of the topographical distribution of neurofibrillary tangles in post-mortem tissue. However, such approaches may not take full advantage of information contained in neuroimaging data. The present study employs an unsupervised data-driven method to identify spatial patterns of tau-PET distribution, and to compare these patterns to previously published “pathology-driven” ROIs. Tau-PET patterns were identified from a discovery sample comprised of 123 normal controls and patients with mild cognitive impairment or Alzheimer’s disease (AD) dementia from the Swedish BioFINDER cohort, who underwent [^18^F]AV1451 PET scanning. Associations with cognition were tested in a separate sample of 90 individuals from ADNI. BioFINDER [^18^F]AV1451 images were entered into a robust voxelwise stable clustering algorithm, which resulted in five clusters. Mean [^18^F]AV1451 uptake in the data-driven clusters, and in 35 previously published pathology-driven ROIs, was extracted from ADNI [^18^F]AV1451 scans. We performed linear models comparing [^18^F]AV1451 signal across all 40 ROIs to tests of global cognition and episodic memory, adjusting for age, sex and education. Two data-driven ROIs consistently demonstrated the strongest or near-strongest effect sizes across all cognitive tests. Inputting all regions plus demographics into a feature selection routine resulted in selection of two ROIs (one data-driven, one pathology-driven) and education, which together explained 28% of the variance of a global cognitive composite score. Our findings suggest that [^18^F]AV1451-PET data naturally clusters into spatial patterns that are biologically meaningful and that may offer advantages as clinical tools.

## 1. INTRODUCTION

Alzheimer’s disease (AD) is neuropathologically defined by the presence of widespread extracellular plaques containing amyloid-β and intracellular neurofibrillary tangles consisting of aggregated tau proteins [Braak and Braak, 1991; Masters et al., 1985]. While amyloid-β may be present decades prior to symptom onset [Jansen et al., 2015], the presence of neocortical tau is temporally more closely related to current cognitive status and degree of neurodegeneration, as convincingly demonstrated by studies utilizing post-mortem tissue, animal models, cerebrospinal fluid and, more recently, the positron emission tomography (PET) tracer [^18^F]AV1451 [Arriagada et al., 1992; Bejanin et al., 2017; Cho et al., 2017; Nelson P. T. et al, 2013; Ossenkoppele et al., 2016; Van Rossum et al., 2012]. [^18^F]AV1451 binds paired helical filaments of tau with high affinity and selectivity [Chien et al., 2013; Lowe et al., 2016; Marquié et al., 2015; Marquié et al., 2017; Xia et al., 2013], and can be used to investigate the distribution of tau pathology in the living human brain. Several studies have shown strong spatial resemblance between *in vivo* tau PET patterns and neuropathological staging of neurofibrillary tangles as proposed by Braak and Braak [Cho et al., 2016; Schöll et al., 2016; Schwarz et al., 2016], reflecting prototypical progression from (trans)entorhinal (stage I/II) to limbic (stage III/IV) to isocortical (stage V/VI) regions [Braak and Braak, 1991]. Furthermore, regional [^18^F]AV1451 retention co-localizes with sites of brain atrophy or hypometabolism [Ossenkoppele et al., 2016; Xia et al., 2017] and has been associated with impairments in specific cognitive domains [Bejanin et al., 2017; Cho et al., 2017; Ossenkoppele et al., 2016].

Given this strong regional specificity of tau pathology, it is important to consider how regions-of-interest (ROIs) are defined, as they could potentially impact study outcomes. To date, most studies employing tau-PET tracers involved ROIs constructed based on neuropathological studies. For example, some studies mimicked the Braak stages *in* vivo [Cho et al., 2016; Schöll et al., 2016; Schwarz et al., 2016], while others selected specific regions reflecting early (e.g. entorhinal cortex) or more advanced (e.g. inferior temporal cortex) disease stages [Johnson et al., 2016]. These approaches have several advantages as they are supported by fundamental research and enhance generalizability across studies. However, compared to neuroimaging, neuropathological data typically include only a few slices in a constrained number of brain regions, and brain tissue is affected by death [Scheltens and Rockwood, 2011]. Additionally, tau PET signal does not equal presence of tau pathology. There are several sources of [^18^F]AV1451 signal and noise, including target binding, off-target binding (e.g. Monamine oxidase, neuromelanin, vascular lesions, iron), non-specific binding and imaging related noise (e.g. partial volume effects) [Choi et al., 2017; Harada et al., 2018; Ikonomovic et al., 2016; Lockhart et al., 2017; Lowe et al., 2016; Marquié et al., 2015; Ng et al., 2017; Schöll et al., 2016]. An alternative approach could therefore be to select ROIs based on data-driven approaches [Dickerson et al., 2011; Grothe et al., 2017; Landau et al., 2011; Pankov et al., 2016], thereby taking full advantage of the abundance of information contained in neuroimaging data, but also accounting for the idiosyncrasies of PET imaging data.

In light of ongoing efforts to define appropriate ROIs and determine tau PET-positivity, it is important to compare data-driven approaches (agnostic, “where is the tau?”) with theory-derived ROIs based on post-mortem studies (directed, “is the tau here?”). In the present study, we applied an unsupervised algorithm to identify clusters of [^18^F]AV1451 signal and compared the spatial patterns of these clusters with neuropathologically derived ROIs described in previous publications. As a secondary analysis, we tested which ROIs best correlated with global cognition in an independent cohort of cognitively normal, mild cognitive impairment and AD dementia subjects. We hypothesized that our data-driven approach would corroborate neuropathological findings, but would also present novel information leading to enhanced associations with cognition.

## 2. MATERIALS AND METHODS

### 2.1 Participants

Two separate cohorts were included in this study. Participants from the Swedish BioFINDER study were used to perform clustering analysis on [^18^F]AV1451 data, whereas participants from the Alzheimer’s Disease Neuroimaging Initiative (ADNI) were used to test associations between the clustering-derived ROIs and cognition. This design allowed us to not only probe the patterns of spatial covariance of [^18^F]AV1451, but also to assess these utility of these patterns as a general [^18^F]AV1451 biomarker without concern of overfitting or “double-dipping” (c.f. [Kriegeskorte et al., 2009]).

Demographic, clinical and biomarker information for both cohorts are presented in Table 1.

**Table 1:** Demographic information, MMSE scores and amyloid-positivity rates

|  | Controls |  | MCI |  | AD |  | Total |  |
| --- | --- | --- | --- | --- | --- | --- | --- | --- |
|  | BioF | ADNI | BioF | ADNI | BioF | ADNI | BioF | ADNI |
| **n** | 55 | 43 | 21 | 37 | 47 | 10 | 123 | 90 |
| **Age**<br>**(SD)** | **75.0**<br>**(6.2)** | **70.3**<br>**(5.9)** | 70.8<br>(10.9) | 72.0<br>(6.8) | 70.1<br>(8.6) | 73.3<br>(4.3) | 72.4<br>(8.4) | 71.3<br>(6.1) |
| **% Male** | 50.9% | 46.5% | 57.1% | 67.6% | 55.3% | 60.0% | 53.7% | 56.7% |
| **Education**<br>**(SD)** | **12.0**<br>**(3.7)** | **16.1**<br>**(2.4)** | **11.7**<br>**(3.7)** | **16.9**<br>**(2.7)** | **12.2**<br>**(3.2)** | **15.0**<br>**(3.0)** | **12.0**<br>**(3.5)** | **16.3**<br>**(2.6)** |
| **% Amyloid+** | 43.6% | 33.3% | **100%** | **44%** | 100% | 100% | **73.3%** | **44.8%** |
| **MMSE**<br>**(SD)** | 29.1<br>(1.1) | 29.0<br>(1.3) | **25.7**<br>**(2.8)** | **28.4**<br>**(2.0)** | **21.2**<br>**(5.1)** | **25.5**<br>**(5.1)** | **25.5**<br>**(4.9)** | **28.3**<br>**(2.5)** |

- \* **BOLD** text indicates significant difference ( $p < 0.05$ ) between cohorts, as measured by t-test, or Fisher's Exact Tests - ADNI = Alzheimer's Disease Neuroimaging Initiative; BioF = BioFINDER, MMSE = Mini-Mental State Examination; SD = Standard Deviation

The BioFINDER cohort is a multi-site study designed for the purpose of developing biomarkers for neurodegenerative diseases. More information can be found at http://biofinder.se. Study participants included 55 subjects with normal cognition, 21 with mild cognitive impairment (MCI), and 47 with Alzheimer’s dementia, who had complete MRI and [^18^F]AV1451 PET data (Table 1). Patients with MCI were referred to a memory clinic and demonstrated objective cognitive impairment that could not be explained by another condition. AD dementia patients met criteria for the DSM-V [American Psychiatric Association, 2013] and NINCDS-ADRDA [McKhann et al., 2011] for probable AD, established by clinicians blinded to PET data. To optimize overlap with the ADNI cohort, dementia patients were only included if they presented with an amnestic-predominant phenotype. Both dementia and MCI patients were only included in this study if they demonstrated abnormal Aβ1-42 levels in the CSF (INNOTEST, cut-off: 650 ng/l; Palmqvist *et al*., 2015). The sample of controls selected for [^18^F]AV1451 scanning was intentionally enriched for β-amyloid positivity to include people in the preclinical stage of AD (see Table 1). This enrichment was achieved by ensuring that 50% of the cognitively normal participants invited for [^18^F]AV1451 imaging had shown positive PET or CSF β-amyloid measurements at previous visits. PET imaging for the study was approved by the Swedish Medicines and Products Agency and the local Radiation Safety Committee at Skåne University Hospital, Sweden. All participants provided written informed consent according to the Declaration of Helsinki, and ethical approval was given by the Ethics Committee of Lund University, Lund, Sweden.

ADNI is a multi-site open access dataset designed to accelerate the discovery of biomarkers to identify and track AD pathology (adni.loni.usc.edu/). The current study included all ADNI individuals with complete [18F]AV1451 scans that were available in November, 2016. This included 43 cognitively normal elderly controls, 37 patients with MCI, and 10 patients with a recent diagnosis of Alzheimer’s dementia (Table 1).

In addition to imaging data, age, sex, education, diagnosis, amyloid-β status on [18F]florbetapir PET [Landau et al., 2013], and scores from six tests measuring global cognition or activities of daily living were downloaded from the ADNI-LONI website (adni.loni.usc.edu). The cognitive tests were as follows: Mini-Mental State Examination (MMSE) [Folstein et al., 1975]; Clinical Dementia Rating Sum of Boxes (CDRSB) [Hughes et al., 1982]; Alzheimer’s disease Assessment Scale 11 (ADAS11) [Rosen et al., 1984] and 13 (ADAS13) [Mohs et al., 1997]; Everyday Cognition (ECog) [Farias et al., 2008]; Functional Activities Questionnaire (FAQ) [Pfeffer et al., 1982]. We also downloaded the ADNI-MEM score, an episodic memory composite score provided by ADNI [Crane et al., 2012].

### 2.2 Imaging

[^18^F]AV1451 images were processed using separate but nearly identical pipelines across the two cohorts. Acquisition and processing procedures for [^18^F]AV1451 processing in the BioFINDER cohort has been described elsewhere [Hansson et al., 2017]. Scans were reconstructed into 5-min frames and motion corrected using AFNI’s 3dvolreg https://afni.nimh.nih.gov/. Mean [^18^F]AV1451 images were created over a time-window of 80-100 minutes post-injection, and these images were coregistered to each subject’s T1 image in native space. Mean images were then intensity normalized using a complete cerebellar gray reference region to create standard uptake value ratio (SUVR) images. Coregistered MRI images were normalized to the MNI-ICBM152 template using Advanced Normalization Tools (https://stnava.github.io/ANTs/) and the transformation parameters were applied to the SUVR images. Finally, SUVR images were smoothed with an 8mm FWHM Gaussian filter.

For the ADNI cohort, mean 80-100 min [^18^F]AV1451 images, as well as MPRAGE images closest to [^18^F]AV1451 scans, were downloaded from the ADNI-LONI website. Details on acquisition procedures for these [^18^F]AV1451 and MRI images can be found elsewhere (http://adni.loni.usc.edu/methods/documents/). [^18^F]AV1451 images were processed in accordance to procedures described in [Schöll et al., 2016]. Briefly, T1 images were processed using Freesurfer v5.3 and [^18^F]AV1451 images were coregistered to native T1s using Statistical Parametric Mapping 12 (www.fil.ion.ucl.ac.uk/spm/). SUVR images were created using a cerebellar gray reference region and images were normalized to MNI space using the parameters from the coregistered T1. Figure 1 shows mean [^18^F]AV1451 SUVR images stratified by diagnosis and amyloid status for each cohort.

**Figure 1.**
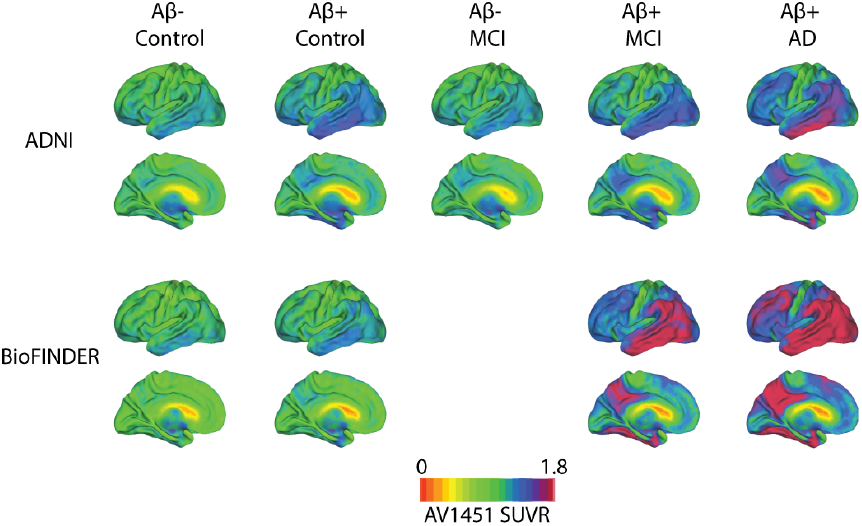
Mean [^18^F]AV1451 uptake according to diagnosis, amyloid status and cohort. Mean [^18^F]AV1451 SUVR images stratified by amyloid status and disease stage, across both the ADNI (top) and BioFINDER (bottom) cohorts.

### 2.3 Clustering of [^18^F]AV1451 data

Our primary analysis involved the derivation of data-driven ROIs by using unsupervised machine learning to elucidate stable patterns of [^18^F]AV1451 signal covariance across a cognitively diverse dataset. Cross-subject [^18^F]AV1451-PET covariance networks were derived from all 123 BioFINDER [^18^F]AV1451 images using an open-source unsupervised consensus-clustering algorithm called Bootstrap Analysis of Stable Clusters (BASC; Figure 2) [Bellec et al., 2010]. BASC is a two-step consensus-clustering algorithm that enhances the stability of the clustering process by repeatedly clustering bootstrapped samples of the input data, and deriving the final partition from this stability matrix, rather than the original data (c.f. [Fred and Jain, 2005]). This approach offers two advantages in the context of this study. First, the stochastic nature of many clustering algorithms tends to lead to different solutions depending on their initialization state, whereas BASC performs clustering on a stability matrix generated from many solutions (and thus many initializations). This leads to greater reproducibility in the clustering solutions generated by BASC. Second, because the initial set of clustering analyses is performed on bootstrap samples of the input data, the final solution is less dependent on the clinical composition of the input data.

**Figure 2.**
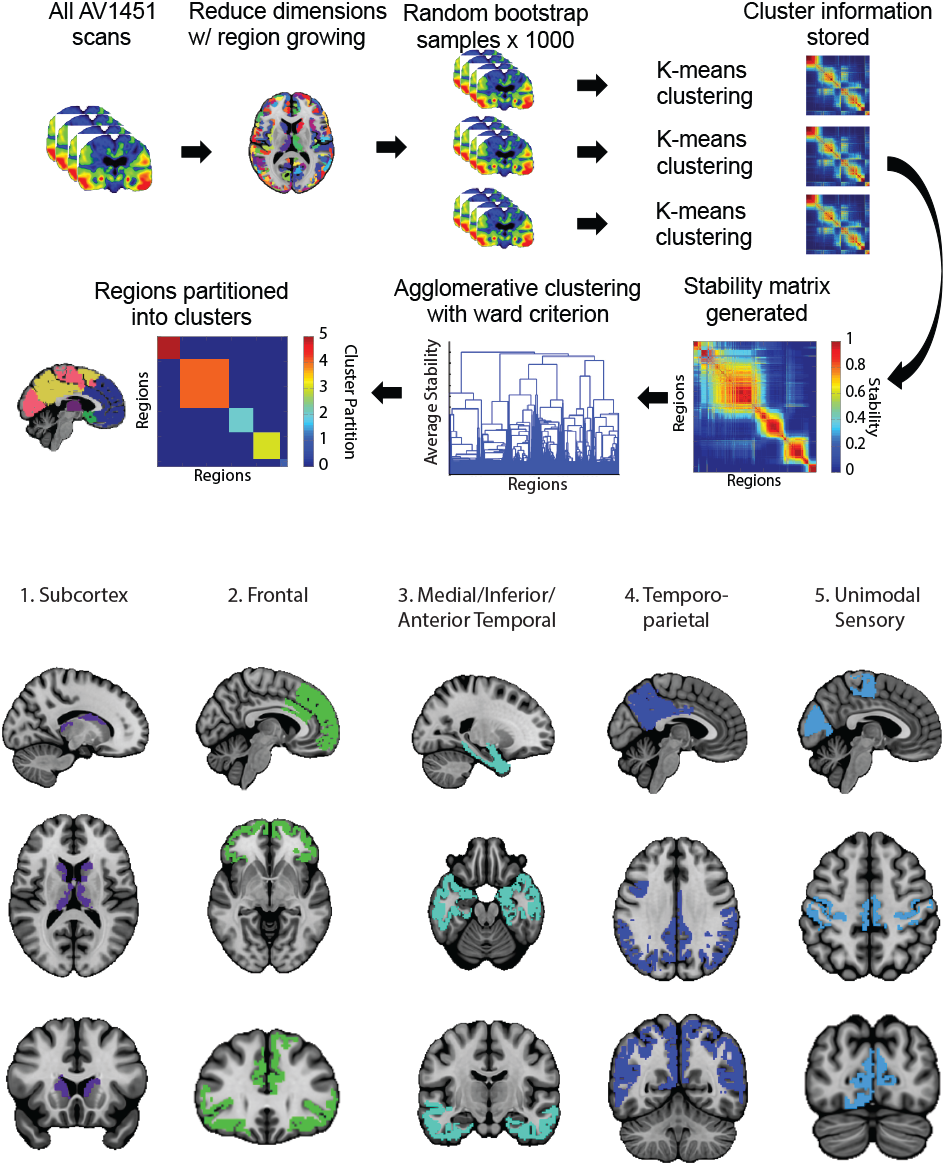
Bootstrap analysis of stable clusters on [^18^F]AV1451 data. [^18^F]AV1451 scans were entered into a voxelwise clustering algorithm. The optimal solutions were determined using the MSTEPS approach. This resulted in five [^18^F]AV1451 covariance networks. These networks were masked with a stability threshold of 0.5, and are displayed in the lower half of the figure.

BASC was adapted to 3D [^18^F]AV1451 data by stacking all 123 BioFINDER [^18^F]AV1451 images along a fourth (subject) dimension, creating a single 4D image to be submitted as input. BASC first reduces the dimensions of the data with a previously described region-growing algorithm [Bellec et al., 2006], which was set to extract spatially constrained atoms (small regions of redundant signal) with a size threshold of 1000mm^3^. In order to reduce computational demands, the Desikan-Killiany atlas [Desikan et al., 2006] was used as a prior for region constraint, and the data was masked with a liberal gray matter mask, which included the subcortex but had the cerebellum manually removed (since this was used as the reference region for [^18^F]AV1451 images). The region-growing algorithm resulted in a total of 730 atoms, which were included in the BASC algorithm. BASC next performs recursive k-means clustering on bootstrapped samples of the input data. After each clustering iteration, information about cluster membership is stored as a binarized adjacency matrix. The adjacency matrices are averaged resulting in a stability matrix representing probabilities of each pair of atoms clustering together (Figure 2). Finally, hierarchical agglomerative clustering with Ward criterion is applied to the stability matrix, resulting in the final clustering solution. The process is repeated over several clustering solutions (k=1 - 50), and the MSTEPs method [Bellec, 2013] was implemented to find the most stable clustering solutions at different resolutions. In the interest of multiple comparisons, and similarity to Braak neuropathological staging (i.e. six ROIs), we chose the lowest resolution solution for subsequent analysis (though the other two solutions are visualized). Note that no size constraints were imposed on clustering solutions (except at the level of atom-size in the region-growing – see above). Cluster-cores were determined as voxels where cluster probability membership exceeded 0.5 (BASC default setting), eliminating unstable voxels from analysis [Bellec et al., 2010; Garcia-Garcia et al., 2018]. After determining cluster-cores in the BIOFINDER cohort, we extracted the average [^18^F]AV1451 SUVR for each cluster core from all ADNI subjects, and these values were used for subsequent analysis investigating associations with cognition.

The choice of the k-means algorithm for the initial clustering and hierarchical clustering with ward criterion for partitioning the stability matrix are somewhat arbitrary. K-means is a particularly fast algorithm and therefore lends itself well to bootstrapping. Meanwhile, the hierarchical clustering routine used in BASC is an appropriate algorithm for the stability matrix, which is a similarity matrix, and it provides solutions at multiple resolutions making it amenable to the BASC framework [Bellec et al., 2010]. Both algorithms are standard, well validated, simple and involve few free parameters. This latter point is important, as BASC itself only has a few principle parameters: namely the number of clusters to extract (in this case, determined by MSTEPS), the number of bootstrap samples (in this case, 500), and the size of the bootstrap sample (in this case, the length of the input data – 123 cases) [Bellec et al., 2010; Orban et al., 2015]. Other parameters are associated with some of the steps peripheral to the central BASC algorithm, namely the region growing preprocessing step and MSTEPS algorithm to determine the number of clusters, and these parameters were left to their default settings. Briefly, the region growing includes a threshold parameter limiting the maximum size of “atoms”, which is mostly related to computational demand. Meanwhile, MSTEPS works on a sparse grid and includes a parameter specifying the percentage of variance maintained (similar to PCA). In addition, MSTEPS allows the definition of the size of the window within which stable clusters are sought [Bellec, 2013].

### 2.4 Definition of Braak stage ROIs described in other studies

A number of studies have created ROIs mirroring the Braak stages described from pathological studies. To test the utility of our data-driven ROIs vis-à-vis those defined in correspondence to the pathological literature, we recreated the Braak ROIs described in three different studies [Cho et al., 2016; Schöll et al., 2016; Schwarz et al., 2016]. Schöll, Lockhart et al. and Cho et al. were constructed using regions from the Desikan-Killiany atlas, and we recreated these ROIs in direct correspondence to what has been reported in these two studies. Schwarz et al. instead generated small ROIs designed to mirror the slabs of cerebral cortex extracted during autopsy for Braak staging. These regions were constructed with a script generously provided by the authors. For all analyses, Braak ROIs were included both individually (“single”) and cumulatively (“stage”). For example, for Braak Stage III, one ROI was created containing all regions from Braak I, II, and III included (“stage”), as well as a ROI created including only regions in Braak III (“single”). Finally, some studies have chosen to use only the bilateral inferior temporal lobe from the Desikan-Killiany atlas to summarize global tau burden [Johnson et al., 2016], so we included this region in subsequent analysis as well. Studies also frequently used the bilateral entorhinal cortex from this atlas, and it should be noted that this region is also included, namely as Stage I from Cho et al. and Schöll, Lockhart et al. Size-weighted average [^18^F]AV1451 SUVR was extracted for each ROI (35 in total) for each subject.

### 2.5 Similarity between data-driven clusters, anatomical ROIs and Braak Stage ROIs

We compiled descriptive information about the similarity between our cluster-derived ROIs and the Braak ROIs from the literature. For comparisons to regions from Schöll, Lockhart et al. and Cho et al., we used normalized mutual information. Due to the small size of the Schwarz et al. regions, comparisons involved measuring the percentage of each Schwarz ROI falling inside of each cluster-derived ROI.

### 2.6 Reproducibility of [^18^F]AV1451 clustering solution

After clustering [^18^F]AV1451 data using BASC (section 2.3), we assessed whether we could reproduce these clusters in a separate dataset. BASC was therefore run on 90 [^18^F]AV1451 scans from ADNI with the exact same parameters used for the BioFINDER dataset. MSTEPS was again used to define the number of clusters. In order to compare the clustering solution to the solution found in the BioFINDER sample, we matched clusters from the ADNI sample to the most spatially similar clusters from the BioFINDER sample, and harmonized the numeric labels between the two solutions. As a qualitative analysis, we extracted voxels that were part of the same cluster in both clustering solutions. The resulting voxels can be thought to represent regions that demonstrated consistent clustering behavior ([^18^F]AV1451 signal covariance) across the two samples. For each cluster, we calculated the Dice coefficient representing within-cluster agreement between the two clustering solutions. We also performed the same analyses constrained within the cluster-cores from the BioFINDER solution, assuming the agreement should be higher within the cores. We also calculated both the adjusted Rand index and adjusted mutual information score (passing the BioFINDER solution as the “true labels”) as a measurement of overall consistency between the two clustering solutions. To put these measurements into context, we performed five 50% splits of the ADNI data and compared clustering solutions between each split. The purpose of this analysis was to identify whether clustering within the ADNI dataset showed greater or less stability compared to the stability between the ADNI and BioFINDER datasets.

### 2.7 Statistical Analysis

Our secondary analyses were aimed to assess the utility and generalizability of our data-driven covariance networks. We performed linear models between these covariance networks and the scores from six different available test scores assessing global cognition and function (see Table S1). In addition, the scores were summarized using Principal Components Analysis (PCA) using Singular Value Decomposition. The PCA was fit to data from the six cognitive test scores, which were scaled to a 0 mean with unit variance. The first component explained 72% of the total model variance, and was used to transform the cognitive data into a single Global Cognition composite score. For each of the cognitive tests, as well as the composite score, separate general linear models for each ROI (40 in total; our five data-driven clusters and 35 ROIs from the literature) were constructed with cognitive test score as the dependent variable and age, sex and education as covariates. We repeated this analysis for the ADNI-MEM score to test the relationship between [^18^F]AV1451 and episodic memory in all 40 ROIs. Tests surviving Bonferroni correction for multiple comparisons are reported.

In order to identify a sparse set of non-redundant covariates that best describe the global cognitive data in ADNI, we submitted all 40 tau ROIs plus age, sex and education to a Least Absolute Shrinkage and Selection Operator (Lasso) regression-based feature selection routine. The Lasso uses L1 regularization (coordinate descent) to penalize regression coefficients based on their maximum likelihood estimates, and is therefore an optimal approach to select a small number of variables from a large number of collinear covariates. In the current implementation, the degree of penalization is optimized using 10-fold cross-validation. All tau ROIs and demographics were scaled to be mean-centered with unit variance, and entered into the Lasso regression model with the Global Cognition composite score as the dependent variable. Features selected by the Lasso (absolute beta > 0.25) were entered together into a general linear model (GLM) with MMSE as the dependent variable. Additionally, to ensure our results were representative of global cognition and not specific to the composite score, the fitted values from this GLM were used to predict scores of each of the six cognitive tests. Finally, the Lasso was repeated separately for each of the individual test as well.

With the exception of BASC, all statistics were implemented using the pandas, numpy, scipy and scikit-learn [Pedregosa et al., 2012] packages in Python 3.5.2 (https://www.python.org/).

## 3. RESULTS

### 3.1 Participant Characteristics

Table 1 contains demographic information, MMSE scores and amyloid positivity rates for both the ADNI and BioFINDER sample. The sample used for clustering (BioFINDER) demonstrated important differences compared to the sample used for testing (ADNI). BioFINDER subjects were less highly educated across the whole sample, and BioFINDER controls were on average older than ADNI controls. Additionally, the BioFINDER sample demonstrated lower MMSE scores across the whole sample compared to ADNI, including within MCI and dementia groups. Finally, 45% of ADNI subjects were amyloid-positive vs. 73% of BioFINDER subjects, which was primarily related to the fact that only amyloid positive MCI patients were included in the BioFINDER sample.

### 3.2 Data-driven Tau-PET covariance networks

123 BioFINDER [^18^F]AV1451 scans were entered into an advanced clustering algorithm in order to identify networks of regional [^18^F]AV1451 signal covariance across subjects. The MSTEPS algorithm identified five-, nine- and 32-cluster solutions as optimal solutions. The parcellations generated from the three stable clustering solutions are visualized in Supplementary Figure S1. For the purposes of comparing with Braak stage ROIs, we chose the lowest-resolution solution (k=5) for subsequent analyses, visualized in Figure 2. The clusters were interpreted and named as follows: “1: Subcortical”, “2: Frontal”, “3: Medial/Anterior/Inferior Temporal”, “4: Temporo-parietal” and “5: Unimodal Sensory”. Cluster 3 bore resemblance to regions often involved in early tau aggregation and atrophy [Braak and Braak, 1991], while Cluster 4 also appeared similar to regions commonly associated with neurodegeneration in AD [Dickerson et al., 2011; Landau et al., 2011]. Of note, the hippocampus was largely unrepresented in any of the cluster-cores, though some voxels in the head of the hippocampus were included in Cluster 3, and a few distributed voxels were included in Cluster 1 (Subcortex). However, using a winner-takes-all clustering approach, the voxels in the hippocampus were almost equally distributed between Cluster 1 and Cluster 3.

### 3.3 Similarity to Braak ROIs

Descriptive metrics were used to quantify the spatial similarity between the data-driven covariance networks and the Braak Stage ROIs introduced in the literature (Figure 3). Cluster 5 (“Unimodal Sensory”) demonstrated a high degree of overlap with Braak Stage VI across all region sets. Spatial similarity was also evident between Cluster 3 (“Medial/Anterior/Inferior Temporal”) and Stage I-IV from Cho et al., and this cluster almost completely circumscribed Stages I-III from Schwarz et al. Cluster 1 (“Subcortex”) was most similar to Schöll, Lockhart et al. Stage II, due in part to its inclusion of the hippocampus. Little spatial similarity was evident between Cluster 2 (“Frontal”) and any of the Braak Stage ROIs, though some similarity was seen with the Stage V region from Schöll, Lockhart et al. and Cho et al. due to their inclusion of many frontal lobe structures. Similarly, Cluster 4 (“Temporo-parietal”) did not demonstrate strong spatial similarity to any of the Braak ROIs, though it did partially overlap with the Braak single IV and V regions from Schwarz et al.

**Figure 3.**
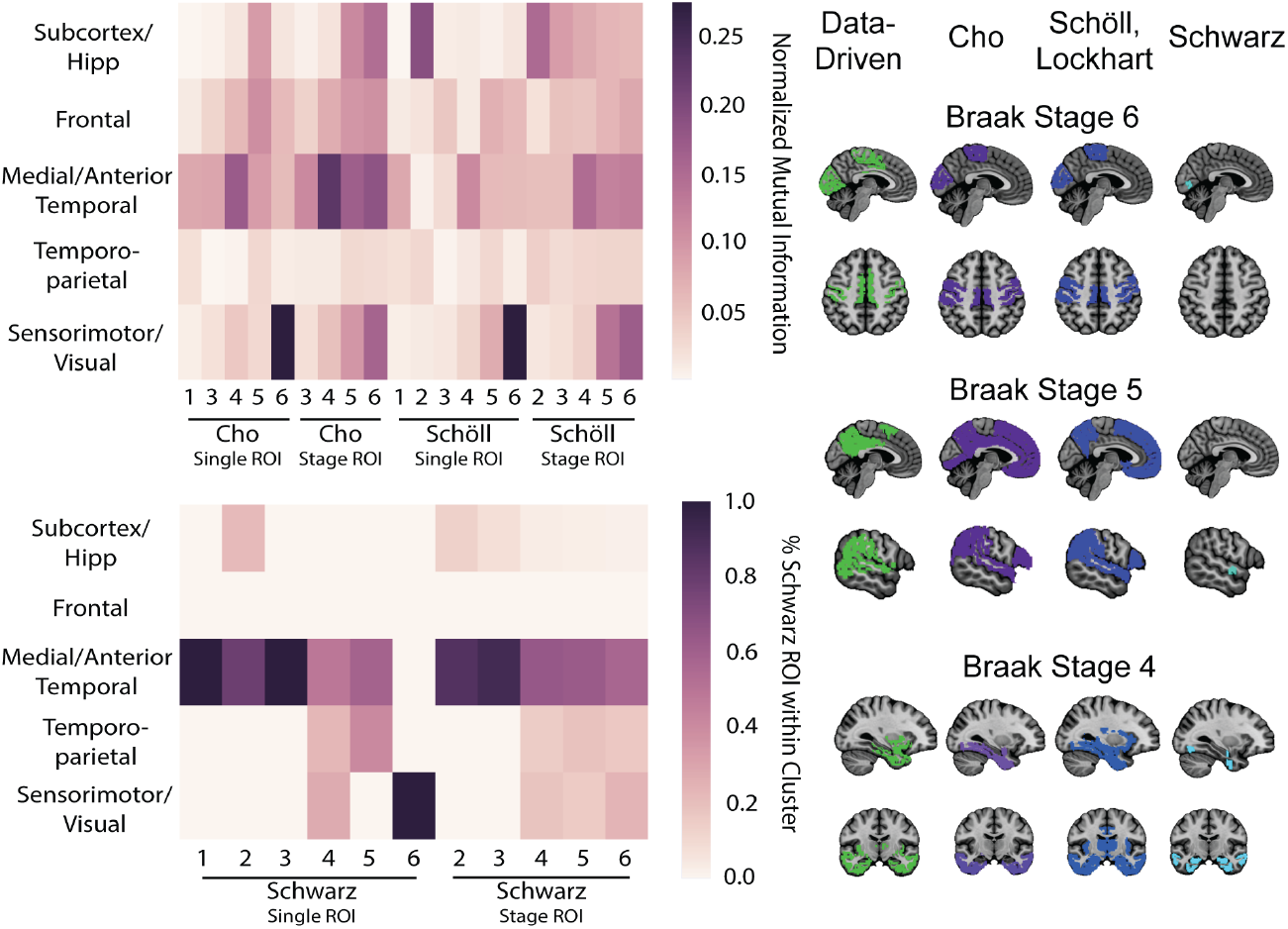
Comparison between data-driven and hypothesis-driven ROIs. Data-driven [^18^F]AV1451 covariance networks were compared to previously existing Braak Stage ROIs from the literature using descriptive statistics. The clusters were compared to ROIs from Schöll, Lockhart et al. and Cho et al using Normalized Mutual Information (top left), and were compared to regions from Schwarz et al. using the percentage of Schwarz ROI voxels within each data-driven cluster.

### 3.4 Associations with cognition in ADNI

General linear models were run in the ADNI dataset assessing associations separately between each of 40 tau ROIs (our five data-driven clusters established in the BioFINDER study, and 35 ROIs from the literature) and a Global Cognitive composite score, controlling for age, sex and education (Figure 4). [^18^F]AV1451 signal in several ROIs demonstrated strong associations with global cognition, though only the data-driven Cluster 4 (“Temporo-parietal”; β = −3.24 [SE=0.91], t = −3.43, p<0.001) survived multiple comparisons.

**Figure 4.**
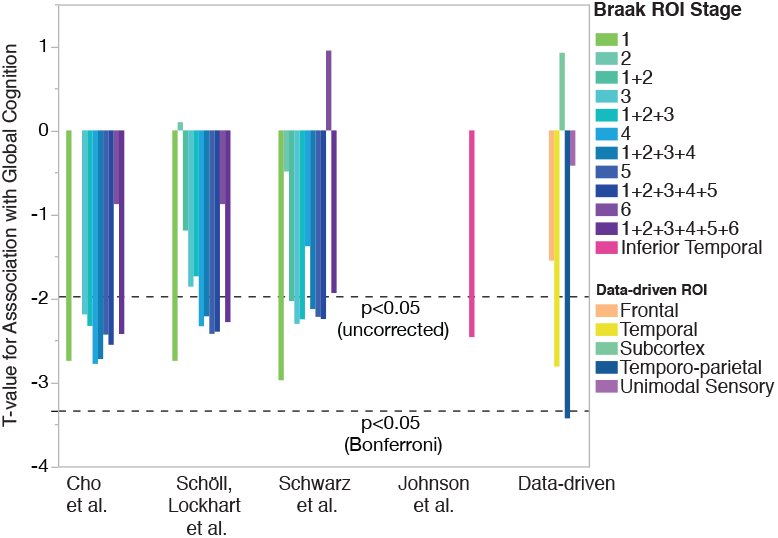
Associations between [^18^F]AV1451 ROIs and global cognition. General linear models comparing [^18^F]AV1451 signal to Global Cognition composite scores were run, adjusting for age, sex and education. For each model, a different [^18^F]AV1451 ROI was used. ROIs included the five clusters identified in our analysis, as well as Braak stage regions taken from three different papers: Schöll, Lockhart et al., 2016 *Neuron*; Cho et al., 2016 *Ann. Neurol*.; Schwarz et al., 2016 *Brain*. Two versions of each Braak ROI were created, one using regions from that stage only (e.g. Stage 3), and one combining all regions from that stage with all regions from previous stages (e.g. Stage 1+2+3). The effect size (t-value) of each tau ROI is shown. [^18^F]AV1451 binding in several ROIs demonstrated strong relationships with Global Cognition, though only the data-driven Temporo-parietal region survived multiple comparisons.

To ensure our results were not specific to the Global Cognition composite score, we repeated this analysis using the six individual measures of global cognition and function that composed the composite score (Table S1). The data-driven Cluster 4 (“Temporo-parietal”) described global cognition better than all other ROIs using four of the six cognitive measures, and was in the top five for all of them. Across all cognitive measures, Clusters 4 and 3 (“Medial/anterior/inferior temporal”) ranked best and second best, respectively, at describing global cognitive data (Figure 5). Notably, the Schwarz Stage I ROI also performed well across cognitive measures, except for the MMSE.

**Figure 5.**
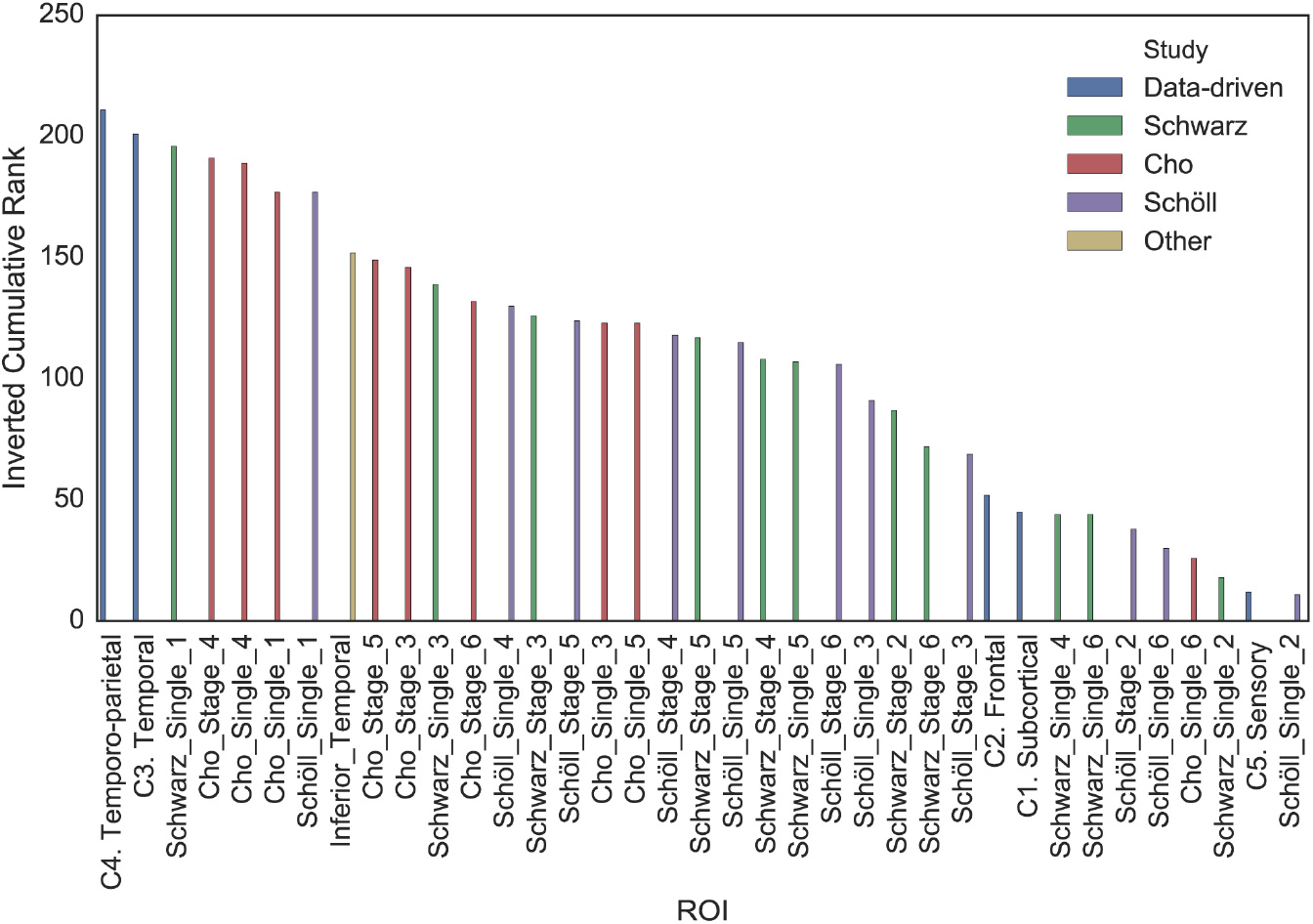
Cumulative ranking of ROI performance across all measures of global cognition and function. For each measure of global cognition, [^18^F]AV1451 ROIs were ranked from worst to best (such that the worst region would have rank of 1) with respect to the effect size of the association between [^18^F]AV1451 in that region and the cognitive score. The ranks were then summed across all cognitive measurements and are displayed here. The data-driven Cluster 4 (“Temporo-parietal”) ranked the best cumulatively across cognitive tests, with the data-driven Cluster 3 (“Medial/Inferior/Anterior temporal”) ranking second best.

Finally, since many ADNI subjects had either MCI or were at early stages of dementia and may not show great variation in tests of global cognition scores, we repeated the above analysis substituting global cognition with a composite measure of episodic memory. (Table S2) shows the top five ROIs with the strongest associations with episodic memory. Although none of the associations survived correction for multiple comparisons, the strongest associations were found with early stage pathological ROIs (resembling (trans)enthorinal cortex), followed by the data-driven temporo-parietal ROI.

### 3.5 Identifying a combinatorial tau-PET biomarker for cognition

Next, all tau ROIs were entered into a Lasso regression model in order to identify a sparse set of covariates that best describe global cognitive data (Figure 6). The optimal penalization value was defined through cross-validation as 0.019. The Lasso reduced all coefficients except Cluster 4 (“Temporo-parietal”), Braak Stage VI from Schwarz et al., and education. These three variables were entered together into a general linear model, and together explained a much greater proportion of variance in global cognitive data (r^2^[4:81] = 0.28, p<0.0001; Figure 6) compared to the individual effect sizes of each covariate (highest r^2^ = 0.12). The earlier negative association between Cluster 4 and Global Cognition was strengthened (t=-4.98, p<0.001), although positive associations were seen for the other two covariates (Schwarz Single 6: t = 3.61, p = 0.001; Education: t = 2.53, p = 0.013). In addition, the fitted values of this GLM explained 18.7 – 26.2% of the variance in the six individual cognitive tests composing the composite score (all p <0.001), indicating the model generalizes well to individual cognitive tests (Table S3). Finally, the Lasso feature selection analysis was repeated for the six individual tests of global cognition. The data-driven Cluster 4 was selected across all six analyses, and was the only ROI selected for two analyses (Table S4).

**Figure 6.**
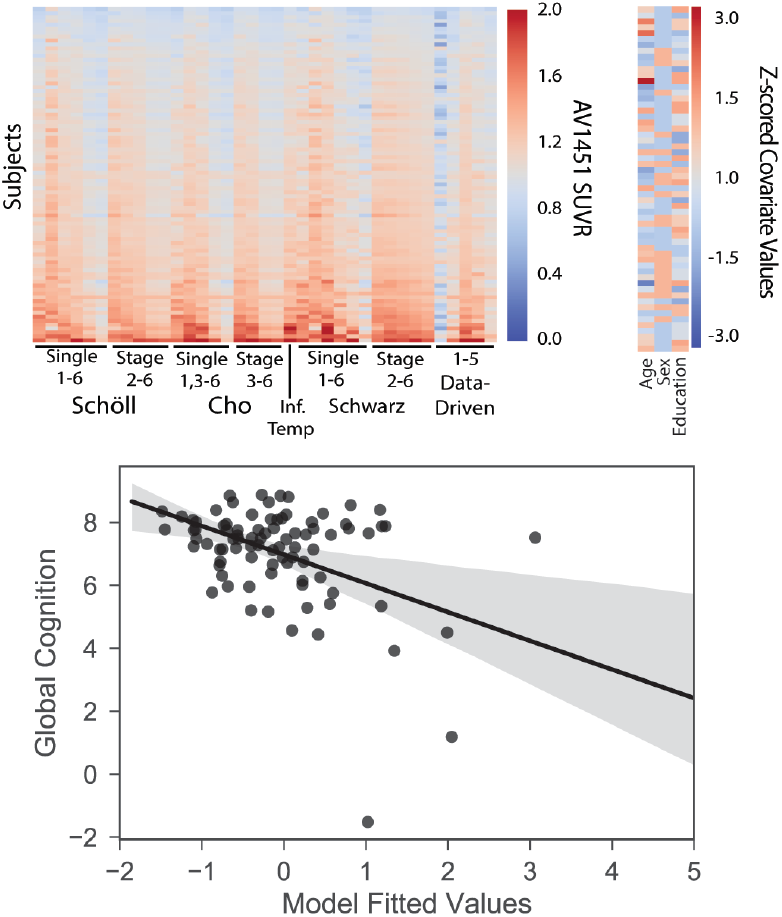
Lasso regression selects most important features related to cognition. All [^18^F]AV1451 ROIs plus age, sex and education were entered into a L1-penalized Lasso regression feature selection routine with the Global Cognitive composite score as the dependent variable. The Lasso selected education and two ROIs: the data-driven Temporo-parietal region, and the Schwarz Single VI region. Together in a general linear model, these features explained 28% of the variance in the Global Cognition score.

### 3.6 Reproducibility of tau-PET clusters across datasets

BASC analysis was run a second time on the 90 ADNI [^18^F]AV1451 scans to establish whether patterns of tau-PET covariance are reproducible across different datasets. MSTEPS identified a six-cluster solution as the lowest resolution solution in the ADNI dataset. Five of these clusters demonstrated similar spatial patterns to the five clusters identified in the BioFINDER sample, while a sixth cluster emerged which uniformly encircled the entire cerebral cortex (Figure S2). This sixth cluster labeled 18% of brain voxels, and the average within-cluster [^18^F]AV1451 SUVR was 0.88 (SD = 0.16). The cluster most likely represents a partial volume or non tau-related atrophy effect, possibly driven by the high proportion of amyloid-negative MCI subjects or the low number of subjects with extensive isocortical tau in the ADNI cohort.

Despite the existence of this sixth cluster and the distinct clinical composition of the two datasets, some agreement between the two clustering solutions could be observed (Figure 7). Overall, 35% of brain voxels showed similar clustering patterns between the two datasets (adjusted Rand index = 0.112; adjusted mutual information score = 0.189). Figure 7A shows a cortical projection of voxels demonstrating similar clustering behavior across both datasets. Across datasets, [^18^F]AV1451 spatial covariance was consistent in the medial and inferior temporal lobes, the primary visual cortex, the temporo-parietal cortex, the medial frontal lobe, and most acutely in the subcortex. The subcortex formed its own cluster in both datasets, both including the hippocampus, and overall showed excellent agreement (Dice coefficient = 0.87). The Dice coefficients in the other clusters ranged from 0.33 – 0.46 (Figure 4B), indicating that around one third to one half of voxels within clusters showed agreement between the two datasets. Notable regions of disagreement included the precuneus and posterior cingulate (clustered with the temporal lobes in ADNI), the insula (clustered with the medial frontal lobe in ADNI), the sensorimotor cortex and the lateral frontal lobes (distributed across multiple clusters in ADNI). When restricting the analysis only to voxels contained within the BioFINDER cluster-cores, the agreement between the two datasets improved (Figure 7B). This observation was consistent across all clusters except the temporo-parietal cluster, and provides evidence supporting the notion that voxels that covary stably within datasets may also show more stable covariance across datasets.

**Figure 7.**
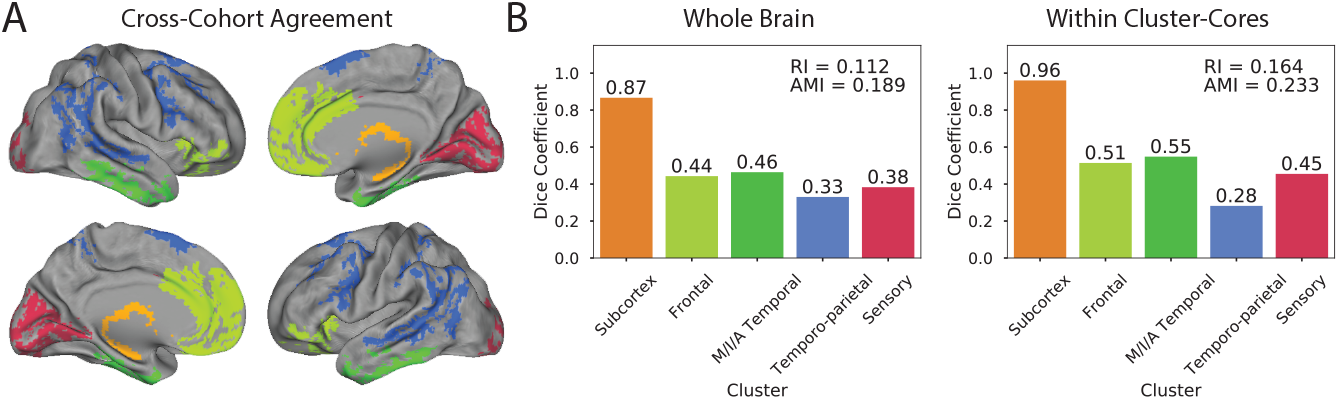
Assessing reproducibility of clusters across cohorts. BASC clustering was performed on ADNI [^18^F]AV1451 data and was compared to the original clustering solution from BioFINDER data. Panel **A**. represents the surface rendering of voxels that shared the same cluster in both BioFINDER and ADNI solutions. Each cluster is represented as a different color. Panel **B**. shows the dice coefficients representing the correspondence between similar clusters in the BioFINDER and ADNI samples. The left graph represents correspondence across the whole brain, while the right graph represents correspondence between clusters within BioFINDER cluster-core masks. RI = adjusted Rand index; AMI = adjusted mutual information score

For the purposes of comparison, BASC was performed on five random 50% splits of the ADNI sample, and the resulting partitions were compared to one another. The average adjusted Rand index across these five within-ADNI train/test splits was 0.166 (SD = 0.031) and the average adjusted mutual information score was 0.225 (SD = 0.021). These within-dataset scores were equivalent to the between-dataset scores when restricted to cluster-cores (adjusted Rand index = 0.164; adjusted mutual information score = 0.233).

## 4. DISCUSSION

In the present study, we applied an advanced unsupervised algorithm to identify clusters of [^18^F]AV1451 signal in 123 subjects ranging from cognitively normal to AD dementia in the Swedish BioFINDER study. Our approach yielded clusters in the temporoparietal, medial/inferior/anterior temporal, unimodal sensory and frontal cortex, as well as the subcortex. In an independent sample of 90 subjects (ADNI), we performed general linear models between tests of global cognition and each [^18^F]AV1451 cluster, adjusting for age, sex and education. In addition, we ran similar models using 35 neuropathologically derived ROIs from previous publications [Cho et al., 2016; Johnson et al., 2016; Schöll et al., 2016; Schwarz et al., 2016]. Several ROIs exhibited strong relationships with cognition, though certain data-driven clusters (temporoparietal and medial/inferior/anterior temporal cortex) appeared to perform slightly but consistently better than other ROIs in ADNI. Supporting this notion, the temporoparietal data-driven cluster was among the three most important features (identified by a Lasso regression model) for predicting global cognition scores. Unsupervised clustering of [^18^F]AV1451 PET data thus revealed the data to self-assemble into stable ROIs resembling well described vulnerable regions in AD, some of which actually enhanced description of cognitive data in an independent dataset. This suggests that data-driven approaches to delineate ROIs may improve clinical utility of [^18^F]AV1451 PET data.

The tau-PET covariance networks derived from our clustering approach exhibited a fair degree of overlap with Braak ROIs derived from autopsy studies, thereby demonstrating biological relevance. Particularly, Cluster 3 (“Medial/Anterior/Inferior Temporal”) was reminiscent of regions involved in early tau accumulation, whereas Cluster 5 (“Unimodal Sensory”) demonstrated a high degree of similarity to regions involved only in the latest stages of AD. In contrast, Cluster 4 (“Temporo-parietal”) did not strongly resemble any of the Braak regions, while its pattern, together with the pattern of Cluster 3, spatially overlapped with cortical regions most vulnerable to neurodegeneration in AD [Dickerson et al., 2011; Landau et al., 2011]. Furthermore, signal in the hippocampus was heterogeneous, adding additional evidence that [^18^F]AV1451 signal in this structure should be interpreted with caution [Cho et al., 2016] [Choi et al., 2017; Ikonomovic et al., 2016]. Similarly, our data-driven approach suggested that most (but not all) frontal lobe structures exhibited [^18^F]AV1451 signal patterns unique to the rest of the cortex. This is notable considering the original Braak Stage V aggregates frontal lobe structures with many of the temporo-parietal structures captured in our Cluster 4. Part of the successful description of cognitive data by the data-driven ROI may be due to its isolation from many of these frontal lobe structures, which may be contributing signal less informative to AD progression, particularly in early disease stages. Finally, our data-driven ROIs provide information that may reconcile some differences between existing Braak ROIs. For example, in our study, [^18^F]AV1451 signal in the putamen and insula covaried with other regions involved in early tau accumulation, which was similar to the ROIs described by Schöll, Lockhart et al., but not Cho et al. (see Table S5 for a summary). However, this pattern was not fully reproduced within the ADNI sample, and so the staging of different ROIs may require further study with larger samples.

Despite the clusters being derived from a sample with several important and disease-relevant differences compared to the testing sample, these data-driven ROIs described global cognitive data slightly better than regions derived from autopsy studies. While the improvement over the other regions was subtle, the increasing movement toward the development of biomarkers demands optimization of ROIs to summarize [^18^F]AV1451 signal [Frisoni et al., 2017; Maass et al., 2017; Mishra et al., 2017]. As such, even small improvements are important for studies assessing more subtle effects of cortical tau accumulation and studies seeking optimal biomarkers for multimodal classification or disease progression [Ota et al., 2015]. The improvement observed is likely due to the data-driven nature of the method used for derivation of the clusters. [^18^F]AV1451 may be binding to several off-target agents, such as (neuro)melanin, iron, vascular pathology and MAO-A/B [Choi et al., 2017; Lowe et al., 2016; Marquié et al., 2015; Ng et al., 2017], and as such, [^18^F]AV1451 signal is likely a mix of true tau pathology and other off-target and non-specific signals. Deriving the clusters from a sample representing a wide breadth of disease stages and additionally including subjects unlikely to have significant cortical tau pathology enhances the likelihood of isolating true tau signal, which covaries strongly and in a regionally specific pattern across disease stages. Additionally, deriving the clusters voxelwise allows freedom from anatomical borders, which may impose unnecessary constraints irrelevant to the spread of tau. Finally, despite its many limitations, multi-subject automatic whole-brain sampling is a distinct advantage of [^18^F]AV1451-PET over pathological studies. This advantage may further enhance the efficacy of data-driven approaches to ROI generation, which evaluate regions equally that may otherwise be overlooked.

Still, ROIs based on pathology remain important in understanding relationships between tau burden and cognition. In our study, ROIs representing the earliest stages of tau pathology, especially the entorhinal cortex, showed the strongest association with episodic memory in a cohort of individuals with normal cognition, mild cognitive impairment and early AD dementia. This finding supports previous literature highlighting relationships between medial temporal lobe tau pathology and decline in episodic memory [Maass et al., 2018]. However, it is noteworthy that the data-driven temporo parietal ROI was again among the top performing ROIs in describing episodic memory, despite the absence of medial temporal lobe structures within this ROI.

The results of this study thus suggest a possible advantage of data-driven approaches in evaluating [^18^F]AV1451 PET data as a biomarker for AD. This study adds to a rapidly growing body of data-driven [^18^F]AV1451-PET studies that have helped to characterize features of this tracer in the context of AD. Sepulcre and colleagues employed a similar unsupervised clustering approach on a set of cognitively intact elderly individuals, which, similar to our study, revealed [^18^F]AV1451-PET covariance between regions of early- and later-stage tau accumulation [Sepulcre et al., 2017]. This suggests these patterns of signal covariance are stable even in the earliest disease stages, lending credence to the use of data-driven biomarkers in multiple contexts. Meanwhile, Jones et al. used a data-driven Independent Components Analysis approach to summarize [^18^F]AV1451 data [Jones et al., 2017]. While the authors concluded the resulting ROIs represented functional brain networks, three of the ROIs bore a striking similarity to those generated by our clustering approach. Our approach builds on these previous studies by assessing relationships between data-driven ROIs and cognition, and by comparing them with other existing ROIs. Maass et al. employed a series of *a priori* and supervised data-driven methods to generate [^18^F]AV1451 ROIs and found a relative equivalence between these ROIs in their association with cognition and a number of other disease markers [Maass et al., 2017]. However, consistent with our study, Maass et al. found [^18^F]AV1451 signal to covary most strongly within a specific set of AD vulnerable-regions, and conclude that these regional measures may perform better than whole-brain ROIs, particularly regarding associations with cognition.

The consistencies across these studies are also underscored by the consistent patterns of cross-subject [^18^F]AV1451 spatial covariance found across the two datasets in the current study. Despite the fact that the ADNI cohort had many fewer subjects with extensive tau burden, and despite differences in the demographic and clinical characteristics between the ADNI and BioFINDER cohorts, unsupervised clustering of [^18^F]AV1451 data revealed a level of consistency between these two datasets that rivaled the consistency of clustering within the ADNI dataset alone. Certain patterns of tau-PET accumulation emerged in key regions across both cohorts. However, the patterns of tau-PET covariance were not entirely consistent between the two datasets, which could reflect true heterogeneity across samples, or could be a matter of instability due to the relatively small sample sizes (particularly in ADNI). However, better consistency between datasets was found within the cluster-cores – regions of greatest clustering stability within the BioFINDER dataset. This finding, alongside the performance of these cluster-cores as biomarkers in ADNI, suggests some degree of cluster stability may be achieved with the BASC approach, even with smaller sample sizes.

We employed a widely used feature selection routine to identify those regions most informative in describing association between [^18^F]AV1451 signal and cognitive data. The feature most strongly associated with global cognition was the data-driven temporo-parietal cluster, which harbored a strong negative relationship when included with the other selected features (p<0.001). The feature selection also resulted in the selection of Schwarz et al. Stage VI and education, both of which associated positively with MMSE in a general linear model. The finding of an association between education and MMSE controlling for tau pathology is consistent with the concept of cognitive reserve [Stern, 2012], and suggests that more highly educated subjects may experience preserved cognition in the face of tau pathology [Hoenig et al., 2017]. While the selection of Schwarz Stage VI is less obvious, possible explanations include partial volume effects and age-related off-target or non-specific signal. Because very few ADNI subjects demonstrate strong [^18^F]AV1451 signal in this ROI, higher [^18^F]AV1451 signal may be related to the presence of more cortex (and thus more off-target or non-specific binding) rather than increased tau pathology. Similarly, off-target [^18^F]AV1451 signal in the cortex and subcortex has been shown to increase with age [Choi et al., 2017; Schöll et al., 2016; Smith et al., 2016], possibly representing binding to reactive astrocytes [Harada et al., 2018] or iron deposits [Choi et al., 2017]. Since age was not selected by the Lasso and therefore was not included in the multivariate model, this may explain the positive association between these regions and global cognition when accounting for [^18^F]AV1451 signal in the temporoparietal region. However, the fact that these ROIs were selected instead of age suggests they may carry additional cognition-relevant information, which may demand further exploration. Regardless, the negative relationship between Cluster 4 (“Temporo-parietal”) and global cognition was substantially increased after regressing out these other variables. This suggests that [^18^F]AV1451-cognition relationships may be enhanced by regressing out off-target or non-specific signal sources.

Our study comes with a number of limitations. First, there were several differences in characteristics between the two samples. We decided to use the BioFINDER cohort for clustering given the broad range of both [^18^F]AV1451 uptake (Figure 1) and cognitive scores (Table 1). As a consequence, our secondary (cognitive) analysis was performed in subjects from the ADNI cohort with more restricted [^18^F]AV1451 uptake and cognitive scores. On a related note, our cluster and results could be influenced by the composition of our samples. However, voxels are only included in the clusters derived for our analysis if the clustering occurs across >50% of bootstrap samples, so it is unlikely that the clustering solution would be strongly driven by, for example, the high proportion of late-stage (i.e. AD) subjects in the BioFINDER sample. Third, contrary to other studies, we did not make an attempt to classify individuals according to stages of tau pathology. Finally, we chose not to apply partial volume correction on our data. Investigating the impact of such corrections is certainly important, but we were interested in the natural behavior of tau-PET signal before any corrections.

In order to aid future studies, we have made the [^18^F]AV1451 clusters from this study available on FigShare (doi = 10.6084/m9.figshare.5758374).

## Acknowledgements

Work at the authors’ research centers was supported by the European Research Council, the Swedish Research Council, the Strategic Research Area MultiPark (Multidisciplinary Research in Parkinson’s disease) at Lund University, the Swedish Brain Foundation, the Swedish Alzheimer Association, the Marianne and Marcus Wallenberg Foundation, the Skåne University Hospital Foundation, and the Swedish federal government under the ALF agreement. This research was additionally funded by Marie Curie FP7 International Outgoing Fellowship [628812] (to R.O.); The donors of [Alzheimer’s Disease Research], a program of BrightFocus Foundation (to R.O.); Author JWV was additionally funded by an Alzheimer Nederland grant and a Vanier Canada Graduate Studies Doctoral award. The authors would like to thank Emma Wolters, Tessa Timmers, Colin Groot, Angela Tam, Alle Meije Wink, Anita van Loenhoud and Elena Kochova for advice and support. The authors would additionally like to thank Adam Schwarz for providing scripts to recreate the ROIs from Schwarz et al., 2016, and Chul Hyoung Lyoo for providing information about creating ROIs from Cho et al., 2016. AVID Radiopharmaceutical generously provided the precursor of AV-1451. Data collection and sharing for this project was funded in part by the Alzheimer’s Disease Neuroimaging Initiative (ADNI) (National Institutes of Health Grant U01 AG024904) and DOD ADNI (Department of Defense award number W81XWH-12-2-0012). ADNI is funded by the National Institute on Aging, the National Institute of Biomedical Imaging and Bioengineering, and through generous contributions from the following: AbbVie, Alzheimer’s Association; Alzheimer’s Drug Discovery Foundation; Araclon Biotech; BioClinica, Inc.; Biogen; Bristol-Myers Squibb Company; CereSpir, Inc.; Eisai Inc.; Elan Pharmaceuticals, Inc.; Eli Lilly and Company; EuroImmun; F. Hoffmann-La Roche Ltd and its affiliated company Genentech, Inc.; Fujirebio; GE Healthcare; IXICO Ltd.; Janssen Alzheimer Immunotherapy Research & Development, LLC.; Johnson & Johnson Pharmaceutical Research & Development LLC.; Lumosity; Lundbeck; Merck & Co., Inc.; Meso Scale Diagnostics, LLC.; NeuroRx Research; Neurotrack Technologies; Novartis Pharmaceuticals Corporation; Pfizer Inc.; Piramal Imaging; Servier; Takeda Pharmaceutical Company; and Transition Therapeutics. The Canadian Institutes of Health Research is providing funds to support ADNI clinical sites in Canada. Private sector contributions are facilitated by the Foundation for the National Institutes of Health (www.fnih.org). The grantee organization is the Northern California Institute for Research and Education, and the study is coordinated by the Alzheimer’s Disease Cooperative Study at the University of California, San Diego. ADNI data are disseminated by the Laboratory for Neuro Imaging at the University of Southern California

## Author Contributions

J.W.V. and R.O. conceptualized and designed the study. P.S., A.C.E., O.H. and R.O. supervised the study. J.W.V., N.M., Y.I.M., T.O.S., M.S., P.B., O.H. and R.O. acquired, processed and analyzed the data. J.W.V., W.F. and R.O. drafted the manuscript. All authors provide critical or conceptual support and revised the manuscript.

## SUPPLEMENTARY

**Table S1.** Best-ranking [^18^F]AV1451 ROIs at describing global cognition across different cognitive tests

| MMSE |  |  |  | CDRSB |  |  |
| --- | --- | --- | --- | --- | --- | --- |
| Rank | Study | ROI | t | Study | ROI | t |
| 1 | Data-driven | Temporo-parietal | -2.45* | Schwarz | Stage I | 3.60** |
| 2 | Data-driven | Temporal | -2.00* | Data-driven | Temporo-parietal | 3.49** |
| 3 | Cho | Single IV | -1.97 | Cho/Scholl | Stage I | 3.49** |
| 4 | Cho | Stage IV | -1.83 | Data-driven | Temporal | 3.33** |
| 5 | Other | Inferior Temporal | -1.80 | Cho | Stage IV | 3.18* |

| ADAS11 |  |  |  | ADAS13 |  |  |
| --- | --- | --- | --- | --- | --- | --- |
| Rank | Study | ROI | t | Study | ROI | t |
| 1 | Data-driven | Temporo-parietal | 4.10** | Data-driven | Temporo-parietal | 3.02* |
| 2 | Schwarz | Stage I | 3.50** | Schwarz | Stage I | 2.57* |
| 3 | Cho | Single IV | 3.43** | Data-Driven | Temporal | 2.35* |
| 4 | Data-driven | Temporal | 3.39** | Cho/Scholl | Stage I | 2.34* |
| 5 | Cho | Stage IV | 3.33** | Cho | Single IV | 2.34* |

| ECOG |  |  |  | FAQ |  |  |
| --- | --- | --- | --- | --- | --- | --- |
| Rank | Study | ROI | t | Study | ROI | t |
| 1 | Data-driven | Temporo-parietal | 3.68** | Schwarz | Stage I | 2.91* |
| 2 | Cho | Stage IV | 3.48** | Data-driven | Temporal | 2.69* |
| 3 | Schwarz | Stage I | 3.40** | Cho/Scholl | Stage I | 2.68* |
| 4 | Cho | Single IV | 3.40** | Data-driven | Temporo-parietal | 2.67* |
| 5 | Data-Driven | Temporal | 3.40** | Cho | Stage III | 2.63* |

\* p<0.05 \*\* p[Bonf.]<0.05
ROI = Region of Interest; MMSE = Mini-Mental State Examination; CDRSB = Clinical Dementia Rating Sum of Boxes; ADAS = Alzheimer's disease Assessment Scale; ECOG = Everyday Cognition; FAQ = Functional Activities Questionnaire

**Table S2.** Best-ranking [^18^F]AV1451 ROIs at describing episodic memory

| ADNI_MEM |  |  |  |
| --- | --- | --- | --- |
| Rank | Study | ROI | t |
| 1 | Schwarz | Stage I | -3.27* |
| 2 | Cho/Schöll | Stage I | -2.99* |
| 3 | Data-driven | Temporo-parietal | -2.85* |
| 4 | Schwarz | Stage II | -2.58* |
| 5 | Cho | Stage III | -2.54* |

\* p<0.05, \*\* p[Bonf.]<0.05

**Table S3.** Fitted values from the General Linear Model comparing selected [^18^F]AV1451 ROIs to Global Cognition composite also explains variance in individual cognitive tests.

| Test | R<sup>2</sup> |
| --- | --- |
| CDRSB | 0.214 |
| ADAS11 | 0.261 |
| ADAS13 | 0.215 |
| FAQ | 0.221 |
| ECog | 0.196 |
| MMSE | 0.219 |

MMSE = Mini-Mental State Examination; CDRSB = Clinical Dementia Rating Sum of Boxes; ADAS = Alzheimer's disease Assessment Scale; ECog = Everyday Cognition; FAQ = Functional Activities Questionnaire

**Table S4.** Variables selected by Lasso regression that optimally described global cognition across different cognitive tests

| MMSE |  | CDRSB |  | ADAS11 |  |
| --- | --- | --- | --- | --- | --- |
| Study | ROI | Study | ROI | Study | ROI |
| Data-driven | Temporo-parietal | Data-driven | Temporo-parietal | Data-driven | Temporo-parietal |
| Data-driven | Subcortical |  |  | Schwarz | Single VI |
| Schwarz | Stage VI |  |  | Schwarz | Stage I |
| Demographic | Education |  |  | Demographic | Education |
| ADAS13 |  | ECOG |  | FAQ |  |
| Study | ROI | Study | ROI | Study | ROI |
| Data-driven | Temporo-parietal | Data-driven | Temporo-parietal | Data-driven | Temporo-parietal |
| Schwarz | Single VI |  |  | Schwarz | Single VI |
| Schwarz | Stage I |  |  | Schwarz | Stage I |
| Schöll | Single II |  |  | Demographic | Education |
| Demographic | Education |  |  |  |  |
| Demographic | Age |  |  |  |  |

MMSE = Mini-Mental State Examination; CDRSB = Clinical Dementia Rating Sum of Boxes; ADAS = Alzheimer’s disease Assessment Scale; ECog = Everyday Cognition; FAQ = Functional Activities Questionnaire

**Table S5.** Disparities in Braak stage regions-of-interests across studies

| Brain Region | Schöll,<br>Lockhart et<br>al. | Cho et al. | Schwarz et al. | Data-driven |
| --- | --- | --- | --- | --- |
| Hippocampus | Included | Not included | Head only | Head only |
| Lingual Gyrus | Stage 3 | Stage 5 | Stage 6 | TP/US |
| Thalamus | Stage 4 | Not included | Not Included | SCN |
| Putamen | Stage 5 | Not included | Not Included | MAIT |
| Lateral Occipital | Stage 5 | Stage 5 | Stage 4 | TP/US |
| PCC | Stage 4 | Stage 5 | Not Included | TP |
| Insula | Stage 4 | Stage 5 | Not Included | MAIT |
| Frontal Lobe | All stage 5 | All stage 5 | Not Included | F/TP |

TP = Temporo-parietal; US = Unimodal Sensory; SCN = Subcortical/Noise; MAIT = Medial/Anterior/Inferior Temporal; F = Frontal

## Supplementary Figures

**Figure S1:**
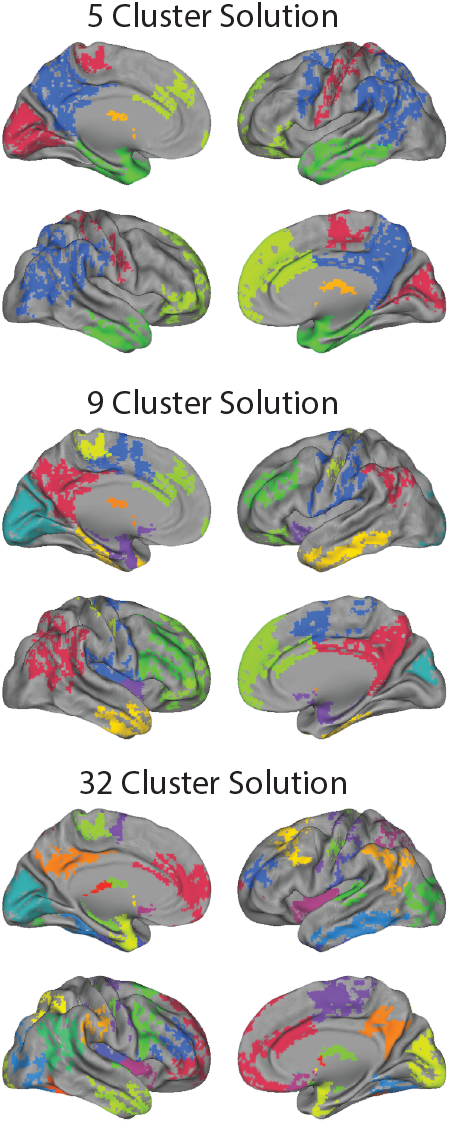
BASC was run on 123 [^18^F]AV1451 images from the BioFINDER cohort. MSTEPS suggested three different resolutions (k=5, k=9 and k=32) to capture the stable patterns of covariance across multiple resolutions. Cluster-core maps were created by setting voxels with cluster stability <0.5 to 0. The cluster-cores from these three solutions are projected onto a cortical surface.

**Figure S2:**
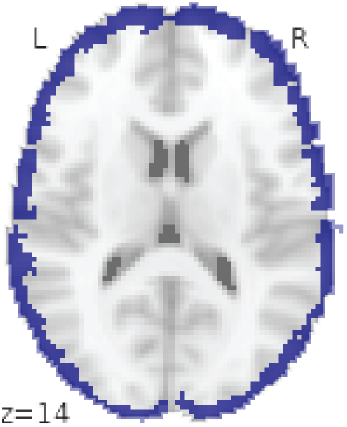
When running BASC in ADNI, a cluster emerged that uniformly surrounded the cerebral cortex, likely representing partial volume effects that could be driven by cortical atrophy in older, amyloid-negative subjects.

